# Range Expansion and Breeding of White-cheeked Duck (*Anas bahamensis*) in the High Andes

**DOI:** 10.1101/2022.07.17.499834

**Authors:** Diego F. Cisneros-Heredia, Mirjaya Izurieta, Emilia Peñaherrera, Maartje Musschenga

## Abstract

We review the distribution of White-cheeked Duck *Anas bahamensis rubrirostris* in mainland Ecuador and show that the species is expanding its range significantly. Contrary to published records, *A. b. rubrirostris* has been present in mainland Ecuador at least since the early 20^th^ century, although probably in low numbers. During the 20th century, the species increased its range along the entire coastlands of Ecuador and nowadays, it has reached the coasts of Colombia. The species has also extended its presence along the Andes, and we report the first breeding records of *A. b. rubrirostris* in the Andes at altitudes between 2360–2440 m, the highest across the entire range of the species. We describe the transitioning plumage between duckling–juveniles, which has not been portrayed in detail before.

---

White-cheeked Duck (*Anas bahamensis*) is widespread but spottily distributed across the Caribbean and South America, where it mainly inhabits brackish waters, mangrove swamps, tidal creeks, estuaries, coastal lagoons, and inland freshwater wetlands, including reservoirs and sewage ponds (Kear 2005, Erize et al. 2006, Johnsgard 2010, Carboneras and Kirwan 2020). Until the 20th century, highland regular records of *A. bahamensis* were only known at Lake Alalay, Bolivia (2550 m), and accidental reports at 3700 m at Lake Poopó, Bolivia, and at 4080 m in Junín, Peru (Bond and Meyer de Schauensee 1943; Fjeldså 1985; Fjeldså and Krabbe 1990). Since the late 20^th^ century, the species has started to disperse into the Andean highlands of Colombia, Ecuador, and Peru (Schulenberg et al. 2007; Freile et al. 2013; Astudillo et al. 2015; Freile et al. 2019a; Rodrıguez-Villamil and Álvarez-Moya 2020).

Three subspecies of *Anas bahamensis* are currently recognized: *A. b. bahamensis* inhabiting the Caribbean and northern Atlantic coasts of South America, south to Brazil; *A. b. galapagensis*, endemic to the Galapagos Archipelago; and *A. b. rubrirostris* from the Pacific coasts and Southern Cone of South America (Kear 2005; Johnsgard 2010; Carboneras and Kirwan 2020). Most information on the breeding biology of *A. bahamensis* is based on data from *A. b. bahamensis* (Sorenson 1992; Sorenson et al. 1992; Kear 2005; Johnsgard 2010; Davis et al. 2017). Accounts about the southern *Anas b. rubrirostris* report that nesting occurs from April–September along the coasts of Ecuador (Marchant 1958, 1960; RSOLAB7 2020), October–November in Argentina (Kear 2005), and November–February in Chile (Saratscheff et al. 1991; Tala and Gabella 1991; Vilina 1995; Rubio C. 1998).

Knowledge on the distribution and breeding of many Ecuadorian waterbird species is scarce, and breeding data has usually been inferred from studies from other areas. Herein, we review the distribution range of *Anas bahamensis rubrirostris* in Ecuador and report the first breeding records in the high Andes of South America.

## METHODS

We made field observations during citizen science activities run by AvesQuito, a citizen collective that promotes bird watching and urban bird ecology studies, and research projects of Universidad San Francisco de Quito USFQ. We have periodically birdwatched since 2010 at the Cumbayá Reservoir, Quito Metropolitan District, province of Pichincha, Ecuador (−0.19483º, -78.42912º, 2360 m) and since 2014 at the Guangopolo Reservoir, Quito Metropolitan District, province of Pichincha, Ecuador (−0.26927° -78.45366°, 2440 m), especially for Quito’s Christmas Bird Count (Cisneros-Heredia et al. 2015). Intensive bird censuses were carried out every two weeks between April and September 2015 at the Cumbayá Reservoir and between 2019 and 2020 at the Guangopolo Reservoir.

We obtained occurrence data from mainland Ecuador from different sources. Published records were synthesized based on a literature review, not limited by study type, study design, or language, conducted in Google Scholar™ scholarly text search (https://scholar.google.com) by online searches. We gathered relevant references using the search terms ‘*Anas bahamensis*’. Open metadata for all occurrences from mainland Ecuador were downloaded from eBird (https://ebird.org) by Cornell Lab of Ornithology (eBird 2020) and iNaturalist (https://www.inaturalist.org) by California Academy of Science and National Geographic (GBIF 2021). Data search and extraction from all sources were conducted in March 2020 and updated in April 2021. For each occurrence point, we compiled geographic data and all other associated information. Protocol for data curation and mining included validation of localities and duplicate detection. All localities were reviewed and validated individually, and coordinates were amended when incorrectly georeferenced in the source. Geographic records of *Anas bahamensis* from mainland Ecuador used for this paper are available in the Supplementary Material.

## RESULTS

Marchant (1958, 1960) recorded several *Anas bahamensis* (*rubrirostris*) between 1954– 1957 in the Santa Elena Peninsula, south-western Ecuador, the first published record in the country (Ridgely and Greenfield 2001). However, an adult male *A. b. rubrirostris* collected at the Santa Elena Peninsula on 22 December 1933 (by Philip Hershkovitz and deposited at the bird collection of Museum of Zoology, University of Michigan; UMMZ 91899, GBIF 2021) provides evidence that the species has been present on the coast of Ecuador at least since the early 20^th^ century—though probably rare based on the lack of collections by expeditions visiting the area during the late 19^th^ century (Chapman 1926). Interestingly, *A. bahamensis* was the most abundant duck at La Carolina, a late Pleistocene site in the Santa Elena Peninsula (Campbell, Jr. 1976), suggesting that the species’ abundance has fluctuated in the region, most probably due to environmental changes.

During the 20^th^ century, *A. bahamensis rubrirostris* increased its range along the Pacific coastlands of Ecuador, becoming locally common below 50 m and evaluated as a non-threatened subspecies in the country (Ridgely and Greenfield 2001, Santander et al. 2013, Freile and Restall 2018, Freile et al. 2019b, eBird 2020, GBIF 2021). There are few inland records on the western lowlands of Ecuador, mainly on the floodplains and rice paddies between Quevedo and Guayaquil (Fig 1) (eBird 2020, GBIF 2021). Ridgely and Greenfield (2001) reported the northernmost Ecuadorian locality of the species in Atacames, province of Esmeraldas, where it has been observed since the 1990s. Solano-Ugalde et al. (2009) evidenced that the species kept advancing north, observing it just 40 km S from the Colombian border (Fig. 1). *Anas bahamensis* was unknown from the Pacific coasts of Colombia until 2013, when Calderón et al. (2013) reported it from the Mar Agrícola farm, in the department of Nariño, ca. 27 km from the Ecuadorian border. Subsequently, there are records of the species up to Buenaventura, department of Valle del Cauca, Colombia, since 2015 (eBird 2020).

**Figure 1.**
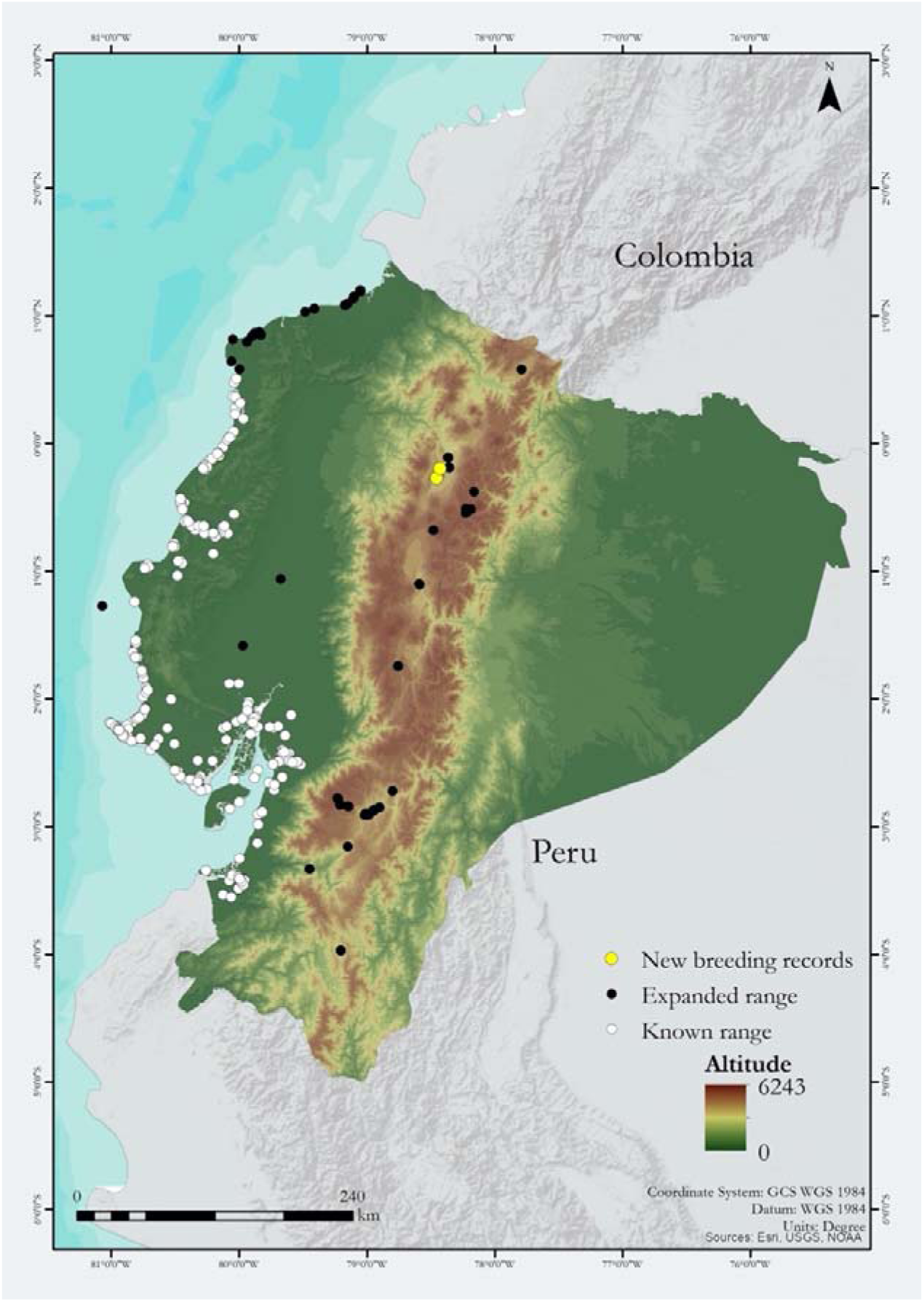
Map of Ecuador showing the distribution range of White-cheeked Duck *Anas bahamensis rubrirostris*. White dots: Records until the late 1990s. Black dots: Records since the early 2000’s, showing the range expansion towards the northern coast, inland western lowlands, and Andean highlands. Yellow dots: New highest breeding records.

The first high altitude reports of *A. bahamensis* in Ecuador was at La Mica Lagoon (3900 m) in 2002 (Lock et al. 2003), where it was recorded again in 2005 and 2006 (eBird 2020, GBIF 2021). Subsequently, it has been recorded at several highland wetlands, including (only first record cited): Cumbaya reservoir (2360 m elevation) in 2009 (Freile et al. 2013); Llaviucu lagoon, Cajas National Park (3160 m) in 2009 (Astudillo et al. 2015); El Paraiso park (2490 m) in 2009 and Ucubamba reservoir (2415 m) in 2012 (Astudillo Webster and Siddons 2013); Quito Airport pond (2350 m) in 2015 (Boyla and Sanchez 2015); Guangopolo reservoir (2440m) in 2017 (Bedoya 2017); Colta lagoon (3310) in 2017 (Morocho 2017); El Salado lagoon (2780 m) in 2017 and Yaguarcocha lagoon (2200 m) in 2018 (Loaiza Bosmediano 2017; Freile et al. 2019a), Jipiro park (2030 m) in 2018 (Hefty 2018), and Yambo lagoon (2600 m) in 2020 (Fattorelli 2020). Records of *Anas bahamensis* across the Andean highlands of Ecuador are becoming more recurrent. The species is present year-round in low numbers at the Cumbaya, Guangopolo and Ucubamba reservoirs and the artificial ponds of El Paraíso Park and Museo Pumapungo (pers. obs.; eBird 2020, GBIF 2021).

On 26 June 2015, a female *A. bahamensis* was observed with eight ducklings swimming in the southern pool of the Cumbaya reservoir (Fig. 2). Ducklings were still covered by down but transitioning to juvenile plumage: face grayish-cream, superciliary band grayish-cream and faint, dark line from eye to nape diffuse, foreneck dark gray, cheeks whitish, lines on sides of back whitish and soft, ventral surfaces whitish with faint lateral stripes, bill bluish gray with light pink wash at the base, eyes brown (Fig. 2). The female and her offspring were either preening or swimming between the northern and southern pools until 06 August 2015. During mid-August, the northern pools were cleaned for sediment removal, and the juveniles were not seen subsequently, but three adults were observed regularly.

**Figure 2.**
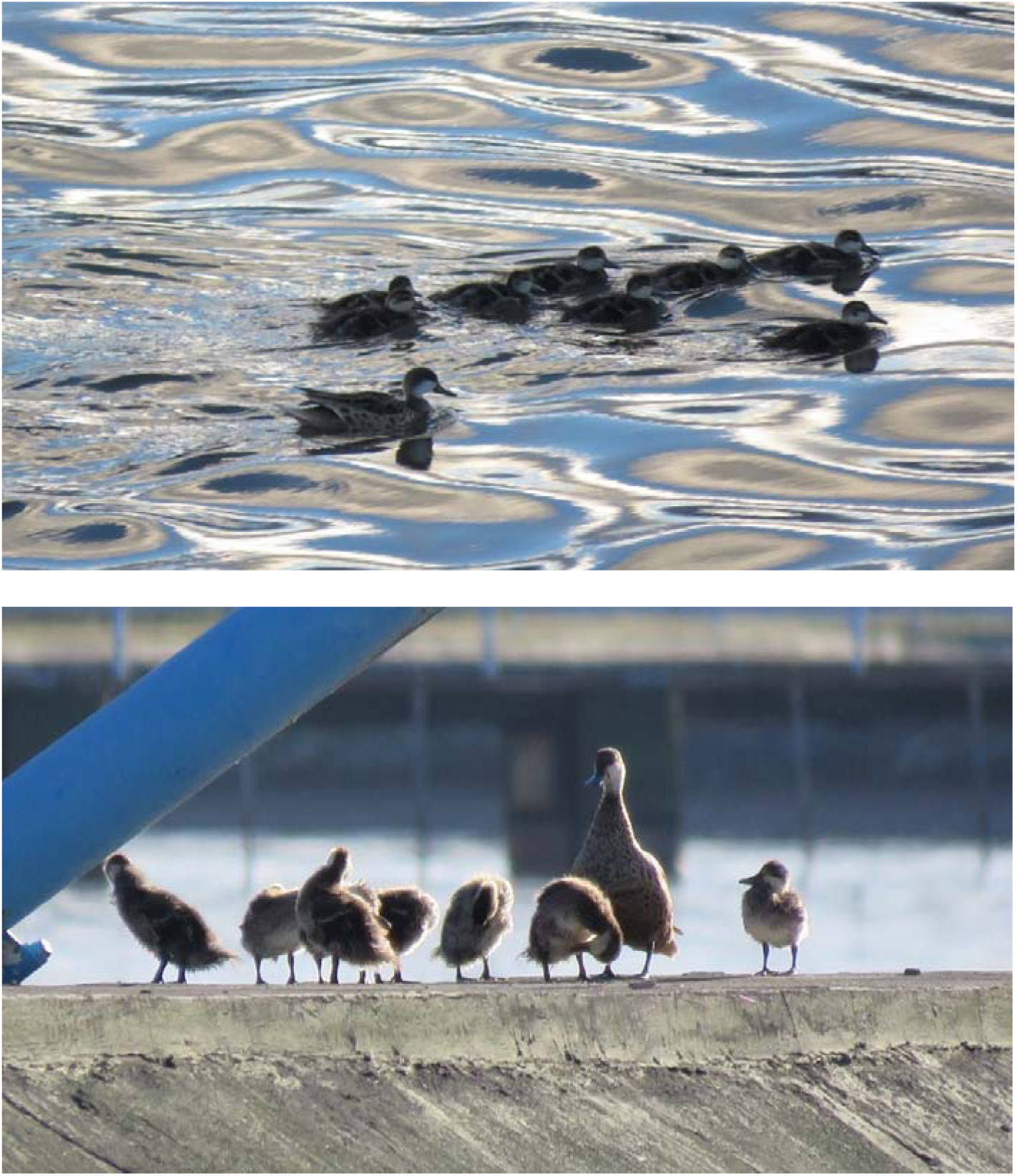
Adult female and eight ducklings of White-cheeked Duck *Anas bahamensis rubrirostris* at the Cumbaya reservoir, Quito Metropolitan District, province of Pichincha, Ecuador, on 26 June 2015.

Between 13–16 November 2019, a female *A. bahamensis* with two ducklings were swimming in a narrow channel, part of the Guangopolo reservoir. On 17 November 2019, two adults were sitting on the channel wall, but there was no trace of the ducklings, and they were not seen subsequently. On 15 July 2020, an adult female with eight ducklings covered by down were swimming in the same channel of the Guangopolo reservoir as in November 2019 (Fig. 3). Duckling plumage is overall the same as described by Carboneras and Kirwan (2020) and Kear (2005). However, lines on the back were yellow, not whitish, especially in the younger ducklings, and there was a brown spot under the eye line, also more visible in the younger ducklings. By 30 July 2020, ducklings were transitioning to juvenile plumage, but the face, cheeks and neck were still yellow, although drabber than in ducklings (Fig. 3). On 04 August 2020, the reservoir was cleaned for sediment removal, and the ducklings were not seen anywhere. On 23 August 2020, the reservoir’s water level was average again, and nine *A. bahamensis* were observed. All ducks had adult size, but bill and plumage were not as bright as in adults suggesting they were juveniles. Their head was smaller and less round, head plumage looked a bit fluffy or downy, and the base of the bill was narrow and ended wider, whereas, in adults, the width of the bill seemed more constant. On 04 September 2020, two juveniles (bill and plumage less bright than adult) were swimming next to each other in the channel. In October 2020, two or three adults and up to five juveniles were in the reservoir (Fig. 3). Subsequently, 8–15 *A. bahamensis* were regular at the Guangopolo reservoir until December 2020, suggesting that the juveniles stayed.

**Figure 3.**
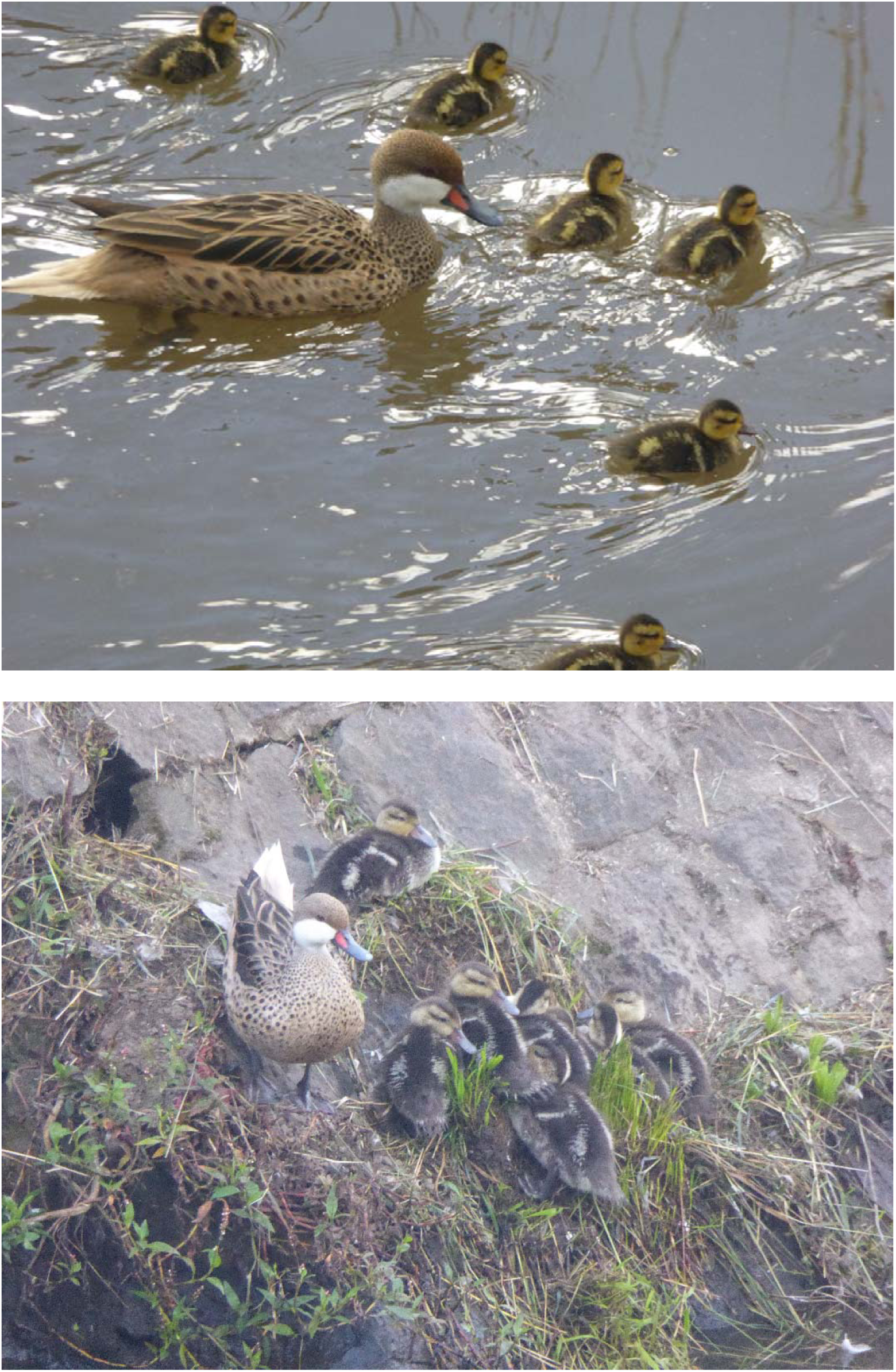
Adult female and eight ducklings of White-cheeked Duck *Anas bahamensis rubrirostris* at the Guangopolo reservoir, Quito Metropolitan District, province of Pichincha, Ecuador, 15 July 2020 (upper) and 30 July 2020 (lower).

**Figure 4.**
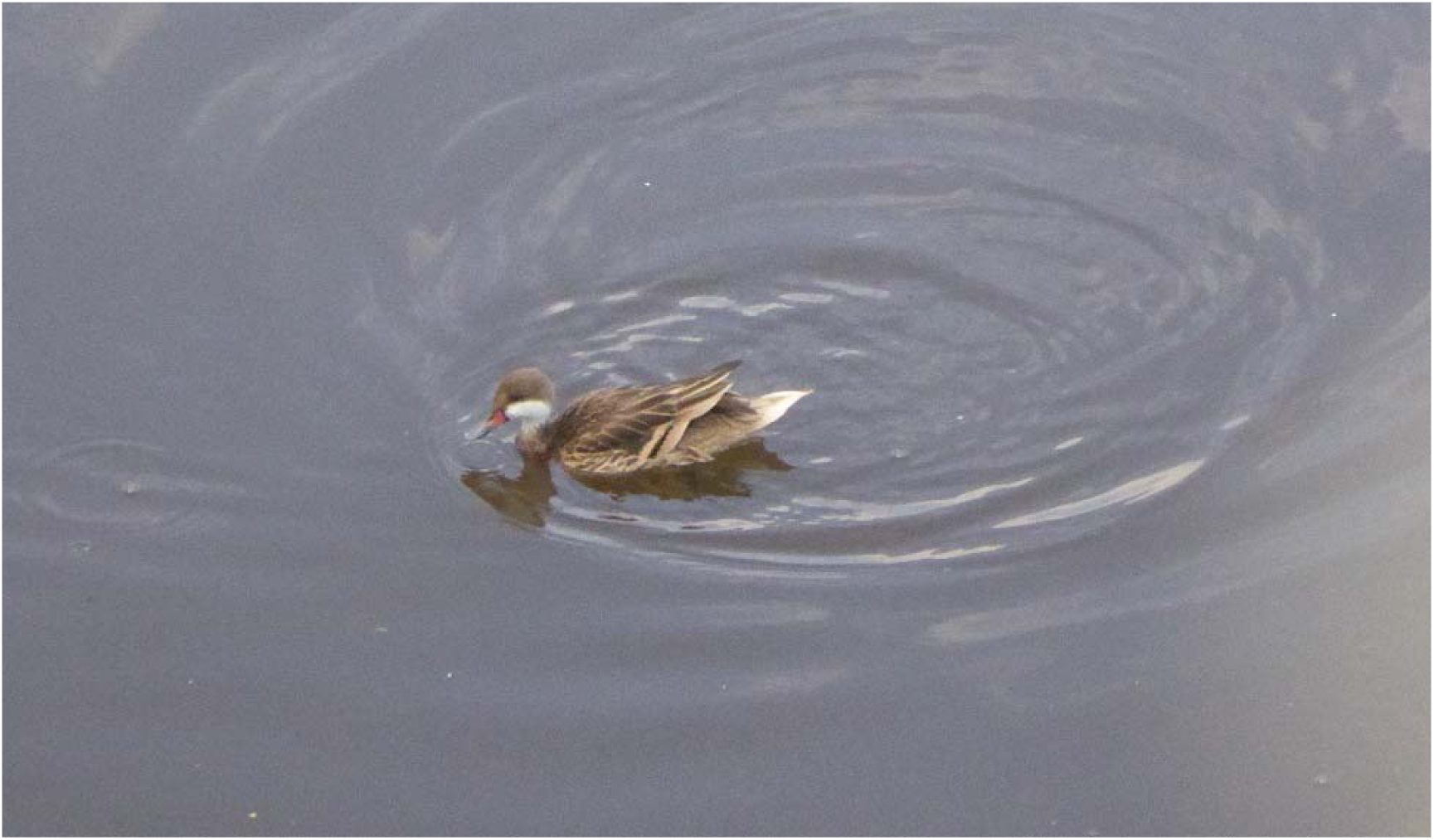
Juvenile of White-cheeked Duck *Anas bahamensis rubrirostris* at the Guangopolo reservoir, Quito Metropolitan District, province of Pichincha, Ecuador, 01 October 2020.

## DISCUSSION

These are the highest breeding records of *A. bahamensis* across its distribution and evidence that the species may be starting to establish self-sustaining populations in the Andean highlands. Reproductive biology was within the reported ranges for brood size, parental care, and fledging time (Kear 2005; Carboneras and Kirwan 2020). Breeding seasonality in the Andes of Ecuador mostly coincides with that reported along the coastlands (April-September), although we also recorded a breeding event in November. All breeding records or long-standing populations in the Andes are in human-made wetlands, probably due to lower impacts by human disturbances, lower predation by domestic and feral dogs and cats (since access to all reservoirs and artificial ponds is controlled), and relatively constant water levels.

## ACKNOWLEDGEMENTS

We would like to thank Empresa Eléctrica Quito EEQ for the permission to access Cumbaya and Guangopolo reservoirs; all members and citizen scientists of AvesQuito for their enthusiastic and constant support; Universidad San Francisco de Quito USFQ, Instituto iBIOTROP, Museo de Zoología and Laboratorio de Zoología Terrestre for logistical and financial support; and xx reviewers for the comments on the manuscript. This paper was possible thanks to the contribution of a wealth of naturalists continuously contributing to eBird and iNaturalist; to scientific collections publishing their curated catalogues in GBIF, and to the Biodiversity Heritage Library for making important literature freely available. This work was supported by Universidad San Francisco de Quito USFQ through research projects (HUBI ID 33 “Diversidad, historia natural, biogeografía y conservación de las aves del Ecuador”, 35 “Estudio de la biodiversidad en áreas urbanas y rurales”, 1057 “Impact of habitat changes on the biological diversity of the northern tropical Andes”, 5452 “Estrés en aves en matrices urbano-rurales en los Andes tropicales”); outreach projects (HUBI ID 278, 292, 483, 607 “Celebrando la Naturaleza: Ciencia ciudadana y educación Ambiental para valorar la biodiversidad”), and operative funds assigned to Instituto iBIOTROP, Museo de Zoología and Laboratorio de Zoología Terrestre.

## Notes

### Competing Interest Statement

The authors have declared no competing interest.

## LITERATURE CITED

Astudillo PX, Tinoco BA, Siddons DC. 2015. The avifauna of Cajas National Park and Mazán Reserve, southern Ecuador, with notes on new records. Cotinga 37: 2–12.

Astudillo Webster P, Siddons DC. 2013. Avifauna de Santa Ana de los Cuatro Ríos de Cuenca. Comisión de Gestión Ambiental de Cuenca, Municipalidad de Cuenca, Universidad del Azuay, Cuenca, Ecuador.

Bedoya J (2017) eBird Checklist - 29 Jan 2017 - Reservorio Guangopolo - 35 species. eBird. https://ebird.org/checklist/S34043296. Accessed 10 Apr 2021

Bond J, Meyer de Schauensee R (1943) The Birds of Bolivia. Part II. Proceedings of the Academy of Natural Sciences of Philadelphia. 95:167–221

Boyla KA, Sanchez M (2015) eBird Checklist - 5 Dec 2015 - Quito airport pond - 16 species. In: eBird. https://ebird.org/checklist/S26138606. Accessed 10 Apr 2021

Calderón JJ, Rosero Y, Ramírez F, et al (2013) Nuevos registros de aves para Nariño y su costa Pacífica. Boletín GAICA 4:5–10

Campbell, Jr. KE (1976) The Late Pleistocene Avifauna of La Carolina, Southwestern Ecuador. Smithsonian Contributions to Paleobiology 27:155–168

Carboneras C, Kirwan GM (2020) White-cheeked Pintail (Anas bahamensis). Birds of the World version 1.0

Chapman FM (Frank M (1926) The distribution of bird-life in Ecuador: a contribution to a study of the origin of Andean bird-life. Bulletin of the AMNH 55: 1–784.

Cisneros-Heredia DF, Amigo X, Arias D, et al (2015) Reporte del 1er Conteo Navideño de Aves de Quito, Ecuador. ACI Avances en Ciencias e Ingenierías 7(2): 37–51. https://doi.org/10.18272/aci.v7i2.256

Davis JB, Vilella FJ, Lancaster JD, et al (2017) White-cheeked Pintail duckling and brood survival across wetland types at Humacao Nature Reserve, Puerto Rico. The Condor 119:308–320. https://doi.org/10.1650/CONDOR-16-169.1

eBird (2020) eBird: An online database of bird distribution and abundance [web application]. Cornell Lab of Ornithology, Ithaca, NY

Erize F, Rodríguez Mata J, Rumboll M (2006) Birds of South America: non-passerines: rheas to woodpeckers. Princeton University Press, Princeton, NJ.

Fattorelli C (2020) eBird Checklist - 30 Jan 2020 - Laguna de Yambo - 28 species. eBird. https://ebird.org/checklist/S63939210. Accessed 10 Apr 2021.

Fjeldså J (1985) Origin, Evolution, and Status of the Avifauna of Andean Wetlands. Ornithological Monographs 36: 85–112.

Fjeldså J, Krabbe N (1990) Birds of the high Andes: a manual to the birds of the temperate zone of the Andes and Patagonia, South America. Zoological Museum, University of Copenhagen, Copenhagen, Denmark.

Freile J, Restall R (2018) Birds of Ecuador. Bloomsbury Publishing, London, UK.

Freile J, Solano-Ugalde A, Brinkhuizen D, et al (2019a) Fourth report of the Committee for Ecuadorian Records in Ornithology (CERO) and a revision of undocumented and erroneous records in the literature. Revista Ecuatoriana de Ornitología 5:52–79. https://doi.org/10.18272/reo.vi5.1277

Freile JF, Ahlman R, Brinkhuizen DM, et al (2013) Rare birds in Ecuador: first annual report of the Committee of Ecuadorian Records in Ornithology (CERO). ACI Avances en Ciencias e Ingenierías (Quito) 5: 24–41. https://doi.org/10.18272/aci.v5i2.135

Freile JF, Santander T, Jiménez-Uzcátegui G, et al (2019b) Lista Roja de las Aves del Ecuador. Ministerio del Ambiente, Aves y Conservación, Comité Ecuatoriano de Registros Ornitológicos, Fundación Charles Darwin, Universidad del Azuay, Red Aves Ecuador, Universidad San Francisco de Quito USFQ, Quito, Ecuador.

GBIF (2021) GBIF Occurrence Download [Anas bahamensis Linnaeus, 1758 Ecuador eBird & iNaturalist]. DOI: 10.15468/DL.WHGPND

Hefty J (2018) eBird Checklist - 9 Dec 2018 - Jipiro Park - 5 species. eBird. https://ebird.org/checklist/S50973338. Accessed 10 Apr 2021

Johnsgard PA (2010) Ducks, geese, and swans of the world. University of Nebraska-Lincoln Libraries, Lincoln, NE.

Kear J (ed) (2005) Ducks, geese, and swans. Oxford University Press, Oxford, UK.

Loaiza Bosmediano JM (2017) eBird Checklist - 9 Jul 2017 - Laguna El Salado - 8 species. eBird. https://ebird.org/checklist/S38069818. Accessed 10 Apr 2021

Marchant S (1958) The birds of the Santa Elena Peninsula, S.W. Ecuador. Ibis 100:349–387. https://doi.org/10.1111/j.1474-919X.1958.tb00404.x

Marchant S (1960) The breeding of some S.W. Ecuadorian birds. Ibis 102(4):584–599

Morocho T (2017) eBird Checklist - 21 Jan 2017 - Laguna de Colta - 15 species. eBird. https://ebird.org/checklist/S34119540. Accessed 10 Apr 2021

Ridgely RS, Greenfield PJ (2001) The birds of Ecuador. Comstock/Cornell Paperbacks, Cornell University Press, Ithaca, NY.

Rodriguez-Villamil DR, Álvarez-Moya WA (2020) Distribución y nuevos registros del Pato Cariblanco (Anas bahamensis) en Colombia. Boletín SAO 29:6–13

RSOLAB7 (2020) White-cheeked Pintail (Anas bahamensis). iNaturalist. https://www.inaturalist.org/observations/60418827. Accessed 10 Apr 2021

Rubio C. M (1998) Nidificación de pato gargantillo (Anas bahamensis) en la Región Metropolitana. Boletín Chileno de Ornitología 5:30–31

Santander T, Ágreda A, Lara A (2013) Censo Neotropical de Aves Acuáticas Ecuador 2008– 2012. Aves y Conservación, Quito, Ecuador.

Saratscheff P, Gabella JP, Tala C (1991) Un breve recorrido por algunos humedales costeros de la V región. Boletín Informativo Unión de Ornitólogos de Chile UNORCH 11:16–17

Schulenberg TS, Stotz DF, Lane DF, et al (2007) Birds of Peru, Revised and updated edition. Princeton University Press, Princeton, NJ.

Solano-Ugalde A, Freile JF, Moscoso P, Prieto-Albuja F (2009) New and confirmative bird records from northern Esmeraldas province, Ecuador. Cotinga 31: 115–118.

Sorenson LG (1992) Variable Mating System of a Sedentary Tropical Duck: The White-Cheeked Pintail (Anas bahamensis bahamensis). The Auk 109:277–292. https://doi.org/10.2307/4088196

Sorenson LG, Woodworth BL, Ruttan LM, McKinney F (1992) Serial monogamy and double brooding in the White-cheeked (Bahama) Pintail Anas bahamensis. Wildfowl 43:156–159

Tala C, Gabella JP. 1991. Observaciones breves. Boletín Informativo Unión de Ornitólogos de Chile UNORCH 12:9–10

Vilina Y (1995) Residencia, abundancia y preferencia de habitat del pato gargantilla (Anas bahamensis) en el humedal “Estero el Yali”, Chile central. Anales del Museo de Historia Natural de Valparaíso 23:89–94.

